# Explain-seq: an end-to-end pipeline from training to interpretation of sequence-based deep learning models

**DOI:** 10.1101/2023.01.23.525250

**Authors:** Nanxiang Zhao, Shuze Wang, Qianhui Huang, Shengcheng Dong, Alan P. Boyle

## Abstract

Interpreting predictive machine learning models to derive biological knowledge is the ultimate goal of developing models in the era of genomic data exploding. Recently, sequence-based deep learning models have greatly outperformed other machine learning models such as SVM in genome-wide prediction tasks. However, deep learning models, which are black-box models, are challenging to interpret their predictions. Here we represented an end-to-end computational pipeline, Explain-seq, to automate the process of developing and interpreting deep learning models in the context of genomics. Explain-seq takes input as genomic sequences and outputs predictive motifs derived from the model trained on sequences. We demonstrated Explain-seq with a public STARR-seq dataset of the A549 human lung cancer cell line released by ENCODE. We found our deep learning model outperformed gkm-SVM model in predicting A549 enhancer activities. By interpreting our well-performed model, we identified 47 TF motifs matched with known TF PWMs, including ZEB1, SP1, YY1, and INSM1. They are associated with epithelial-mesenchymal transition and lung cancer proliferation and metagenesis. In addition, there were motifs that were not matched in the JASPAR database and may be considered as *de novo* enhancer motifs in the A549 cell line.

**Availability:** https://github.com/nsamzhao/Explain-seq

**Supplementary information:** Supplementary data are available as attachment.

## 1 Introduction

Decoding regulatory functions that are encoded in genomic sequences is a major challenge in understanding how genomic variations are associated with phenotypic diseases and traits. High-throughput sequencing methods have been developed to screen for regulatory regions genome-wide. DNase I hypersensitive site sequencing (DNase-seq) (Boyle et al., 2008) is designed to detect genome-wide chromatin accessibility. Transcription factor (TF) binding and histone modifications are measured using Chromatin Immuno-Precipitation sequencing (ChIP-seq) (Johnson, Mortazavi, Myers, & Wold, 2007; Robertson et al., 2007). Enhancers are regulatory DNA sequences that recruit TFs to up-regulate target gene expression in a cell-type specific manner, which governs physiology and development in humans (Arnold et al., 2013). STARR-seq is a massively parallel reporter assay to identify potential enhancers and provide a direct functional or quantitative readout of enhancer activity genome-wide (Arnold et al., 2013). The genome-wide quantitative enhancer map enables interrogating enhancers in higher resolution than binary peak regions such as peaks from DNase-seq.

Deep learning techniques have made substantial progress in modeling of genomic sequence to predict epigenetic marks such as DNA accessibility, TF and histone marks across a range of cell types. Particularly, convolutional neural networks (CNNs), have been shown to accurately predict epigenomic features with DNA sequences (Avsec et al., 2021; Kelley et al., 2018; Zhou & Troyanskaya, 2015). For example, DeepSEA, given any 1000 bp DNA sequence, can accurately predict 919 binary labels for the sequence, representing open chromatin regions, TF binding sites, and histone mark regions altogether in a multi-label CNN model (Zhou & Troyanskaya, 2015). The model significantly outperformed the previous state-of-art gkm-SVM, which leveraged a gapped-kmer-SVM classifier to predict functional sequence elements in regulatory DNA (Ghandi et al., 2016). One potential explanation for why CNN is a better fit for genomic sequence learning tasks is that filters learned during training is analogous to position weight matrices (PWMs) of motifs, which are often conserved and positional invariant.

However, interpreting machine learning models, especially black-box deep learning models, to derive the biological knowledge learned by models remains elusive. Specifically, what sub-sequences make contributions to certain predictions, and how we can summarize those sub-sequences into human-readable motifs? Additionally, due to the limited quantities of conducted STARR-seq experiments, the relationship between enhancer sequences and activities across different cell lines is still poorly understood. There lacks a human enhancer sequence-to-activity model that learns the *cis*-regulatory grammar in a cell-type specific manner, which can accurately predict enhancer activities.

To address these questions, we presented a novel end-to-end pipeline, Explain-seq (**Fig. 1A**), to automate the process of developing and interpreting deep learning models. ENCODE has recently released 6 STARR-seq datasets for common human cell lines (Consortium, 2020). These new datasets provide an opportunity to examine and compare enhancers in a cell-type specific manner. Here we demonstrated Explain-seq by analyzing a new STARR-seq dataset in the A549 lung cancer cell line. The pipeline started with training a CNN with a regression layer at the end of the network to predict cell-line specific enhancer activity and ended with outputting derived predictive motifs. Our trained regression model outperformed the gkm-SVM model in terms of the Pearson correlation coefficient. Also, derived motifs from Explain-seq were matched to other known TF motifs in the JASPAR database including ZEB1, YY1, SP1, and INSM1, which are associated with lung cancer (Fornes et al., 2020). In addition, there were derived motifs not similar to any known motifs, for example, one was enriched with short repeats ATGAAA, which may be considered as *de novo* motifs. The *de novo* motifs discovered by Explain-seq may allow us to design the synthetic enhancers with desired activity in a cell-line specific manner.

**Figure 1.**
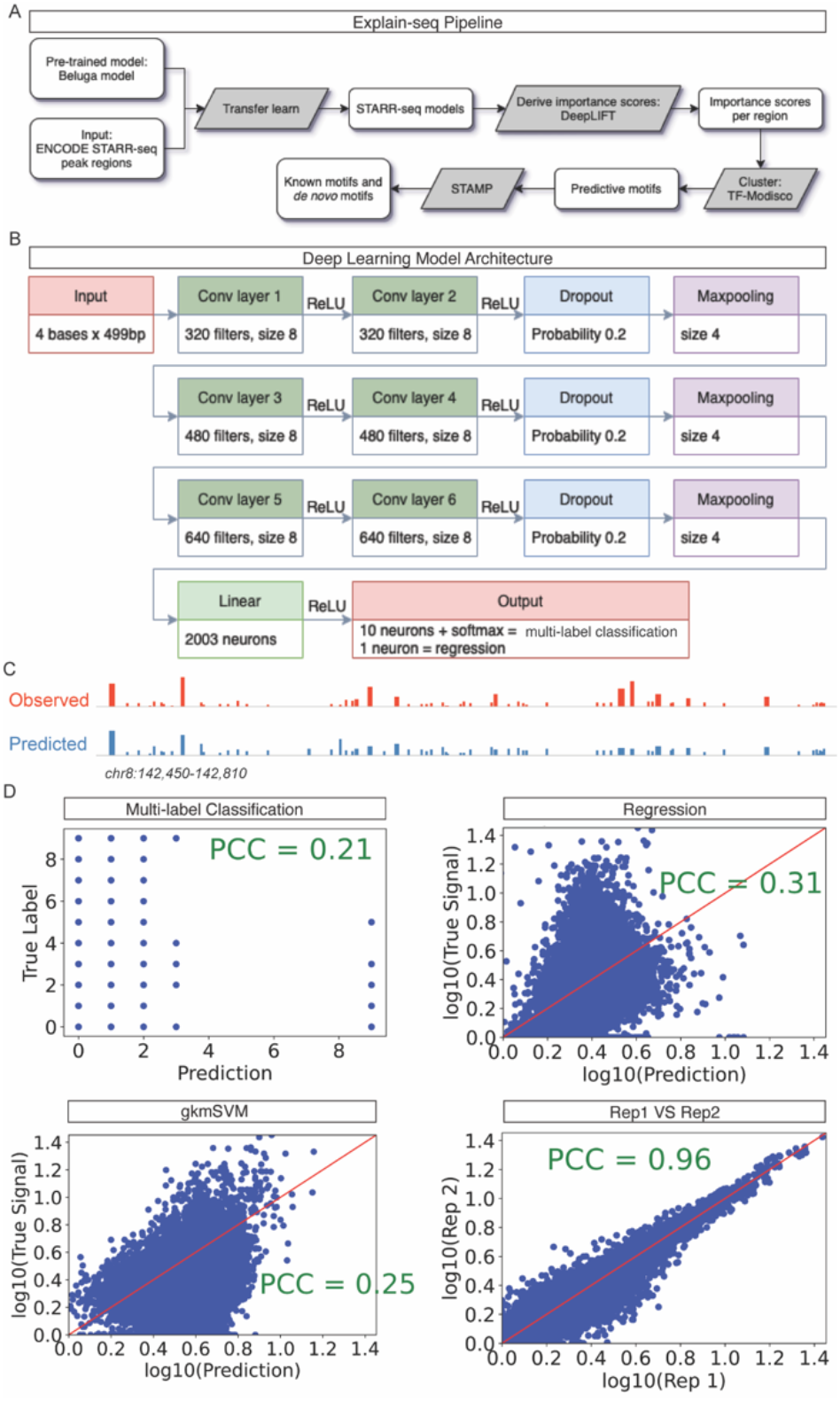
Explain-seq Predicts Enhancer Activity from Genome-wide DNA sequences. (A) Overview of Explain-seq pipeline to infer enhancer activities from the A549 cell line and to identify known and *de novo* motifs. (B) Architecture of the convolutional neural network (CNN) that was trained to predict A549 enhancer activities from 499bp sequences. Both the regression model and the multi-label classification model employ the same architecture. (C) Explain-seq predicts enhancer activity genome-wide. The IGV genome tracks screenshot depicts observed and predicted signals using the regression model for a locus on the held-out test chromosome 8. (D) Explain-seq with CNN regression model predicts enhancer activity better than gkm-SVM and the CNN multi-label classification model. Scatter plots of predicted vs. observed enhancer activity signals across all DNA sequences in the test set chromosomes are shown for CNN multi-label classification model, CNN regression model, and gkm-SVM. We also calculated the correspondence between the actual signals in one biological replicate of A549 with the actual signals in another biological replicate to serve as the maximum prediction accuracy threshold for this task.

## 2 Methods

### 2.1 Data access and preparation

ENCODE phase 4 released 6 STARR-seq datasets on June 11, 2020, which includes common human cell lines MCF-7, HCT116, A549, SH-SY5Y, HepG2, and K562 (Consortium, 2020). We downloaded their peak regions in .bed format and signal values in .bigwig format.

To prepare input data for the deep learning model, we first selected all regions at the summit of each STARR-seq peak and binned them into 499-bp windows. We limited the input sequence length to 499bp since the size selection in STARR-seq experiments before cloning to plasmids is 500 bp (Z. Chen et al., 2021). We included 3 adjacent regions on either side of each selected summit region by sliding a 100 bp overlapping window. We randomly selected 500,000 regions of size 499 bp on the hg38 human genome excluding ENCODE blacklist regions as negative sets (Amemiya, Kundaje, & Boyle, 2019). The signal for each 499-bp window was calculated by the averaging signal value of the whole corresponding region. For the regression model, the average signal values were directly used for training. For the multi-label classification model, we further generated 10 labels using the average value. In total, we have selected 1,027,953 peaks as our input data. We split our input data by chromosome for training, validation, and testing. Specifically, we used chr7 for validation, and chr8 and chr9 were held out for testing.

### 2.2 Overview of Explain-seq pipeline

Explain-seq pipeline is an end-to-end analytical pipeline to discover potential known and *de novo* motifs given genomic sequences (**Fig. 1A**). It takes input as genomic region coordinates with labels for classification tasks (in .bed format) or one-hot encoded sequences with continuous value for regression task (in .mat format) (K. M. Chen, Cofer, Zhou, & Troyanskaya, 2019). In addition, it requires a user-defined deep learning model in PyTorch (Paszke et al., 2019). Optionally, weights from the pre-trained model can be transferred to the new model for transfer learning. After training and validation loss function converges, the model and input sequences are piped into DeepLIFT to compute contribution scores with respect to enhancer activities in a single-nucleotide resolution (Shrikumar, Greenside, & Kundaje, 2017). TF-Modisco is then used to cluster and summarize motifs with the weighted input sequences like weighted by importance scores (Shrikumar et al., 2018). Potential motifs are compared to known motifs in databases through STAMP (Mahony & Benos, 2007). Non-matched motifs are considered as *de novo* predictive motifs in a specific cell line.

### 2.3 Model architecture, training scheme, and transfer learning

We designed our model as a CNN which takes input as one-hot encoded 499-bp long DNA sequences (A = [1,0,0,0], C = [0,1,0,0], G = [0,0,1,0], T = [0,0,0,1]) to predict enhancer activities (**Fig. 1B**). The architecture is inspired by the Beluga model, which has double convolutional layers than DeepSEA, to enable later transfer learning (Zhou et al., 2018). Specifically, our model includes 6 convolutional layers with an equal kernel size of 8 and has 320, 320, 480, 480, 640, and 640 filters for each layer, respectively. ReLU is used as the activation function in each convolutional layer. Every two convolution layers are followed by a dropout layer with a probability of 0.2 and a max pooling layer with a pooling size of 4. Those 6 convolution layers are followed by a fully connected layer with 2003 neurons. We provided two design choices for the final layer. The first option for multi-label classification is 10 nodes where each node represents a label in a specific cell line. For example, the first node representing the probability of the input sequence (or region) has a signal value in [0, 1), and the last node represents in [9, 10). Here, we used binary cross entropy as the loss function. The second option for regression setting is one node for continuous signal value prediction. Mean squared error (MSE) was used as a loss function in regression. The reason for developing and comparing the two designs is that the data might be noisy, and it may not have enough statistical power for us to learn a regression model. Binning the continuous values and learning a multi-label classification model may help denoise the data and learn a more general and representative model.

We implemented our model in PyTorch (Paszke et al., 2019). Considering this is a genome-wide learning task with large data, transfer learning is useful to speed up the training and also to improve accuracy (Avsec et al., 2021; Kelley, Snoek, & Rinn, 2016). We transferred all the weights of convolutional layers from the pre-trained Beluga model as initial weights. The weights from the fully connected layer were excluded since the original Beluga model has an input size of 2,000 bp. After initialization, we fine-tuned the weights by re-training the model. To speed up the development, we used the Selene framework to facilitate the training process (K. M. Chen et al., 2019). We used Adam optimizer with a learning rate of 1e-5 (Kingma & Ba, 2014), a batch size of 128, and used an Nvidia Titan V with 12 GB memory GPU to develop our model.

### 2.4 Baseline comparisons

We compared our CNN-based regression model with another machine learning method gkm-SVM (Ghandi et al., 2016). gkm-SVM trains gapped-kmer SVM classifiers for DNA sequences to detect functional sequence elements in regulatory DNA. We used the Pearson correlation coefficient (PCC) as a metric to compare the prediction accuracy with the true signals. In addition, we calculated the correlation between the actual signals in one biological replicate with the actual signals in another biological replicate to serve as the maximum prediction accuracy threshold for this task.

### 2.5 Derive nucleotide importance scores

DeepLIFT is an algorithm to calculate feature importance scores for neural networks by propagating activation differences (Shrikumar et al., 2017). We used DeepLIFT to derive importance scores for all input with signals larger than 3 with respect to cell-line specific activity. The advantages of DeepLIFT compared to other interpretation methods are: 1) the RevealCancel rule of DeepLIFT allows it to properly handle saturation cases while integrated-based methods may give misleading results (Ancona, Ceolini, Öztireli, & Gross, 2017). 2) DeepLIFT is a good and faster approximation of SHapley Additive exPlanation (SHAP) value (Shrikumar et al., 2017). 3) Di-nucleotide frequency shuffling mimics true genomic background to increase importance scores signal-to-noise ratio. As background, we shuffled 100 times for each input sequence while maintaining di-nucleotide frequency. The output of DeepLIFT is importance scores in nucleotide resolution, and hypothetical importance scores are similar to mutagenesis indicating what importance scores would be placed on a different base in the sequence.

### 2.6 Clustering and summarizing sub-sequences into motifs

TF-Modisco is a clustering-based algorithm to consolidate motifs from sequences with importance scores (Shrikumar et al., 2018). The algorithm started with finding potential motifs, named seqlets, through MEME (Bailey et al., 2009). TF-Modisco implemented a correlation alternative, continuous Jaccard similarity, to better calculate the similarity between seqlets than cross-correlation, and developed the density-adaptive distance to improve clustering on weak motifs when their distances are generally larger (Shrikumar et al., 2018). We specified the final motif size as 15 bp with 5 bp flanking on each side. The target seqlets false discovery rate was set to 0.15 and the max seqlets per cluster was set to 20,000. We sampled 4,900 sequences for our null Laplace null distribution. The output of TF-Modisco contains potential motifs with PWMs, contribution weight matrices with importance scores, and hypothetical score matrices.

### 2.7 Annotation with known motifs

STAMP was leveraged to annotate motifs from TF-Modisco within a known motif database, JASPAR (Fornes et al., 2020; Mahony & Benos, 2007). We trimmed off the motif edges with an information content of less than 0.4 to improve matching accuracy. Since motif-finding procedures from our pipeline or others using importance scores are different from those generated by frequency-based methods, PWM representations varied and were required to be manually compared to known motifs in the final annotation step.

## 3 Results

### Explain-seq predicts enhancer activity from DNA sequence

We designed a computational end-to-end pipeline, Explain-seq, to discover known and potential *de novo* motifs given genomic sequences (**Fig 1A** and **Method**). To learn the *cis*- regulatory code and grammar embedded in enhancer sequences, we have developed a CNN-based deep learning model to predict enhancer activity given DNA sequences (**Fig. 1B**). We have downloaded and pre-processed 6 public STARR-seq datasets in ENCODE Phase 4 (**Method**). The public genome-wide enhancer activity maps provide high-quality datasets to build predictive models of enhancer activity in a cell-line specific manner. To demonstrate Explain-seq usage, we only focused on the A549 cell line to develop our model and pipeline. In total, we had 1,027,953 regions of size 499 bp on the hg38 genome reference for the A549 cell line. Classification-based and regression-based two CNN models have been deployed to map 499 bp DNA sequences to their enhancer activities (**Fig. 1B** and **Method**). The two models’ architectures are similar to that of the Beluga model except for the output layer. The regression model has only one output node in the final layer, while the multi-label classification model has 10 output nodes representing different activity ranges which are then followed by a SoftMax layer to determine the final prediction (**Method**). We transferred all the weights in convolutional layers from the well-trained Beluga model and fine-tuned all parameters when re-trained with STARR-seq datasets (**Fig. 1A**). Transfer learning can speed up training when applied to related task model training, and improve accuracy. Our models’ training and validation loss function converged within 20 epochs given a genome-wide training task (**Supplementary Fig. 1A-D**). After training, we accurately predicted the enhancer activity signals for both models (**Fig. 1C**).

Our regression model outperformed the multi-label classification model and the gkm-SVM model when evaluated on test hold-out chromosomes, chr8 and chr9 (**Fig. 1D**). The predicted and observed enhancer-activity profiles are similar with Pearson correlation coefficient (PCC) 0.31, 0.21, 0.25, for a regression model, multi-label classification model, and gkm-SVM model, respectively. Our regression model outperformed the multi-label classification model which indicates that ENCODE STARR-seq datasets’ signal-to-noise ratios are high enough to be predicted in high resolution with respect to continuous values instead of categorical labels. Even the best-performance regression model is not close to the concordance between experimental replicates (PCC=0.96). It indicates the need for additional information, such as open chromatin and TF binding in the A549 cell line, rather than DNA sequences alone to further boost prediction accuracy. Chen *et al.* trained a model on the same A549 STARR-seq dataset with input as signal shape to predict binary binding (Z. Chen et al., 2021)(Z. Chen et al., 2021). Although they achieved validation AUROC 0.9984 and AUPR 0.9978 using a binary classification model with different preprocessing steps and input, their model was not cell-type specific since the signal shape was transferable between cell lines and was not able to be leveraged for predictive cell-type specific enhancer motifs. We trained a final regression model with both replicates to proceed with interpretation in our pipeline (**Supplement Fig. 1B**, **Fig. 1D**).

### Explain-seq reveals important TF motifs

Interpreting our regression model with Explain-seq reveals known TF motifs in the A549 cell line, and potential *de novo* motifs. Given the regression model, DeepLIFT calculated the importance scores of each nucleotide in the selected input sequence whose mean the signal is larger than 3 (**Fig 2A)**. To summarize all seqlets into readable motifs, TF-Modisco clustered 20,000 seqlets into consolidated potential motifs. Those motifs were then compared to motifs in the JASPAR database using STAMP to verify with known motifs. Briefly, A549 is the most used human non-small cell lung cancer cell line, consisting of hypotriploid alveolar basal epithelial cells. By applying STAMP, we searched for *de novo* motifs and compared them to known PWMs. Overall, we identified 47 known motifs including ZEB1, SP1, YY1, and INSM1 (**Fig. 2B**). ZEB1, zinc-finger e-box binding 1, is part of the ZEB family in humans. ZEB1 is involved in the generation and maintenance of epithelial cell polarity and its expression in epithelial cells results in epithelial-mesenchymal transition (EMT) (Georgakopoulos-Soares, Chartoumpekis, Kyriazopoulou, & Zaravinos, 2020). SP1, specificity protein 1, is important for lung cancer cell proliferation and metastasis during tumorigenesis (Hsu et al., 2012). Transcription factor Yin Yang 1 (YY1) is associated with the EMT process in the A549 cell line and regulates pulmonary fibrotic progression in lung epithelial cells (Zhang et al., 2019). INSM1, identified from STAMP in the A549 cell line, is a sensitive marker for the neuroendocrine differentiation of human lung tumors (Rosenbaum et al., 2015). In summary, the determined known motifs are A549 cell line specific, indicating corresponding biological activities of the hypotriploid alveolar basal epithelial cells from the A549 cell line. Additionally, STAMP enables us to identify *de novo* motifs (**Fig. 2C**). For example, some *de novo* motifs enrich for CpG sites. CpG sites are regions of DNA where a cytosine nucleotide is followed by a guanine nucleotide in the sequence. CpG sites are highly related to DNA methylation that occurs more frequently by hypermethylation in cancers. Given the novel TF motifs, we are able to explore more biological insights within the A549 cell line.

**Figure 2.**
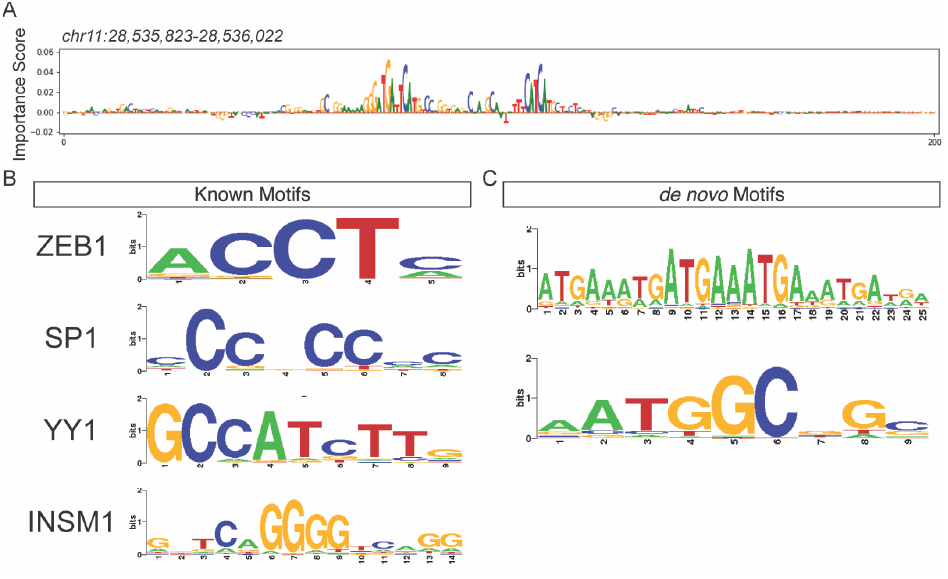
Explain-seq reveals known motifs and *de novo* motifs for enhancer activity. (A) Within Explain-seq, DeepLIFT calculated the importance scores of each base pair in selected input sequences whose mean signal is larger than 3. Important sub-sequences are highlighted. (B) Explain-seq identified known motifs (C) and Explain-seq revealed some *de novo* motifs for the A549 cell line.

## 4 Conclusion

Deep learning algorithms like CNN have been widely used in DNA sequence analysis within the whole genome. However, deep learning model interpretation and biological explanation remain changeling. Here, we introduced a novel end-to-end computational pipeline, Explain-seq, to automate the process of developing and interpreting deep learning models in the context of genomics. We demonstrated the usage with new STARR-seq datasets to predict enhancer activities, characterize *cis*-regulatory code, and identify known and *de novo* motifs using deep learning algorithms. Explain-seq quantitatively predicts enhancer signals in continuous values. Transfer learning is applied to improve the model accuracy and increase computational efficiency. Also, Explain-seq provided insights into the interpretation of the deep learning model and identify biological relevance. Specifically, Explain-seq reveals the sequence-to-function relationship by calculating nucleotide importance scores. Furthermore, by comparing with the existing motif database, Explain-seq successfully determines cell type-specific known and *de novo* motifs, which may contribute to the functionality.

Generalization of predictive models from one cell line to multiple cell lines, and to various cell types of different organisms would be addressed in the future. For example, we anticipate that by integrating various STARR-seq datasets with profound biological interpretation, we are able to ultimately decode gene regulatory information in entire genomes.

## Code and Data Availability

Explain-seq pipeline: https://github.com/nsamzhao/Explain-seq Data at Zenodo: 10.5281/zenodo.7526380

---

**Supplement Figure 1.**
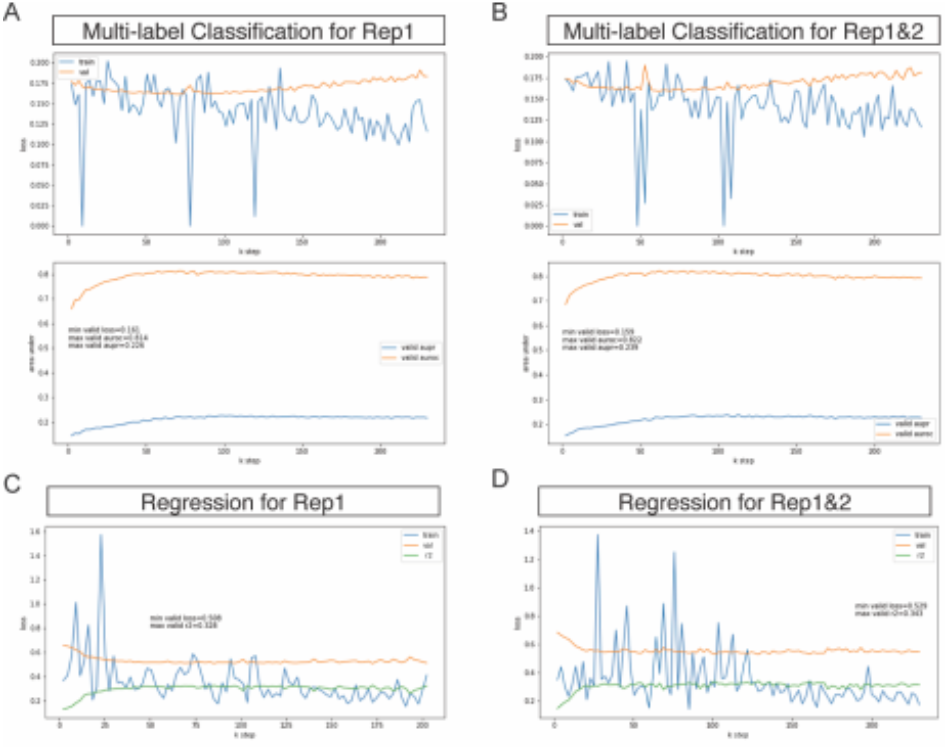
Multi-label Classification and Regression Model Performances. (A) Loss function along the iterations of multi-label classification model for replicate 1 (top row). Model performance on the test data using AUPR and AUROC metrics to measure (second row). (B) Loss function along the iterations of the multi-label classification model for replicate 1 and replicate 2 (top row). Model performance on the test data using AUPR and AUROC metrics (second metrics). (C) Loss function along the iterations of the regression model replicate 1. (D) Loss function along the iterations of the regression model replicate 1 and replicate 2.

